# Harmonized Z-scores calculated from the large scale normal MRI database to evaluate brain atrophy for neurodegenerative disorder

**DOI:** 10.1101/2020.06.09.126748

**Authors:** Norihide Maikusa

## Abstract

Alzheimer’s disease (AD), the most common type of dementia in elderly individuals, slowly and progressively diminishes cognitive function. Mild cognitive impairment is also a significant risk factor to the onset of AD. Magnetic resonance imaging (MRI) images have become widely used to detection and understand the natural progression not only AD but also neurodegenerative disorders. For this purpose, construct a reliable cognitive normal database is important. However, difference in magnetic field strength, sex and age between normal database and evaluation data-set can be affect the accuracy of detection and evaluation of AD and other neurodegenerative disorders.

To solve this problem, we suggest a harmonized Z-score considering differences filed strength, sex and age derived from large cognitive normal subjects dataset ((1235 subjects)) including 1.5 T and 3T T1 brain MRI. And we evaluate our harmonized Z-score of discriminative power of AD, and classification accuracy between stable MCI and progressive MCI.

The harmonized Z-score of hippocampus achieved high accuracy (AUC=0.96) for detection AD and moderate accuracy (AUC=0.70) for classification stable MCI and progressive MCI. Theses results shows that our method not only can detect AD with high accuracy and high generalization capability, but also can be valid to classify stable MCI and progressive MCI.

## Introductions

Alzheimer’s disease (AD) is the most common cause of dementia, and typically shows memory impairment at the earliest clinical stage. Magnetic resonance imaging (MRI) as a biomarker to quantify AD progression has shown much promise. Use of MRI in morphometric or volumetric measurement of brain atrophy, such as changes in cortical thickness, hippocampus volume, whole brain volume, has resulted in improved diagnosis. These measurements can also be used to assess the effectiveness of any applied therapies.

On the other hand, a volume measurement pipeline called FreeSurfer [4,5] has been often used for volume measurement of anatomical regions of interest (ROI) in brain imaging clinical research. However, FreeSurfer need to take 10 to 20 hours to execute, so it is not suitable for clinical applications. Also the single-atlas method that warps a single atlas into the individual brain image has a problem in ROI segmentation accuracy. In this study, we used 30 Japanese brain atlas including 131 anatomical regions for highly accurate segmentation and calculating brain structure volume using the multi-atlas method. We performed segmentation for 1235 normal control (NC) groups (1089 subjects were scanned by 3T MRI and 146 subjects were scanned 1.5 T MRI). And We calculated the harmonized Z-score for each anatomical region of interest from NC group. In harmonization, we consider several confounding factors (i.e. sex, scanner filed strength, 1.5 T or 3T, estimated total intracranial volume (eTIV) and age. After harmonization the brain structure volume, we can minimize the difference between subgroups of each covariates (i.e., the male and female, 1.5 T and 3 T MRI scanner) and evaluate brain atrophy without considering the effects of age and whole brain volume.

## Materials and Methods

### MRI acquisition

The 3three-dimensional (3D) T1-weighted images of the NC were obtained from two different protocols on 3.0 T MRI scanners at National Center of Neurology and Psychiatry: 693 individuals underwent Protocol 1 and the other 438 individuals underwent Protocol 2. On the other hand, all of the patients underwent Protocol 1. Protocol 1: 3.0-T MR system (Philips Medical Systems, Best, The Netherlands) with the following protocol: repetition time (TR)/echo time (TE), 7.18 ms/3.46 ms; flip angle, 10 degree; number of excitations (NEX), 1; 0.68 × 0.68 mm^2^ in plane resolution; 0.6 mm effective slice thickness with no gap; 300 slices; matrix, 384 × 384; field of view (FOV), 261 × 261 mm. Protocol 2: 3.0 T MR system (Verio, Siemens, Erlangen, Germany) with the following protocol: TR/TE, 1800 ms/ 2.25 ms; flip angle, 9 degree; NEX, 1; 0.87 × 0.78 mm^2^ in plane resolution; 0.8 mm effective slice thickness with no gap; 224 slices; matrix, 320 × 280; FOV, 250 × 250 mm. All data were collected after obtaining informed consent from participants and approval from the ethics committee of our center.

We also obtained 1.5T T1-weighted MR images for evaluating dataset from the Japanese Alzheimer’s Disease Neuroimaging Initiative (J-ADNI) data set, provided by National Bioscience Database Center in Japan [2]. Data were acquired using 1.5 T MRI scanners (GE Healthcare, Siemens and Philips) and preprocessed with non-parametric nonuniform normalization (N3) [9] and phantom based distortion correction [8]. We divided the total of 507 participants into four groups, NC, AD, stable MCI (sMCI) and progressive MCI (pMCI) based on criteria as follows:

**NC subjects:** MMSE score 24–30, CDR of 0, non-depressed, no memory complaint.
**AD subjects:** MMSE score of 20–26, CDR 0.5 or 1, and memory complaint.
**sMCI subjects:** MMSE score 24–30, memory complaint (preferably corroborated by an informant), objective memory loss measured, CDR of 0.5, absence of significant levels of impairment in other cognitive domains, and essentially preserved activities of daily living (if the diagnosis was MCI for ≥ 36 months).
**pMCI subjects:** MMSE score 24–30, memory complaint (preferably corroborated by an informant), objective memory loss measured, a CDR of 0.5, absence of significant levels of impairment in other cognitive domains, essentially preserved activities of daily living (if the diagnosis was MCI at baseline but conversion to AD was reported after baseline within 6–36 months).

All of the J-ADNI subjects whose samples were evaluated had taken the MMSE, CDR, ADAS, and FAQ as neuropsychological screening tools. The evaluating MRI data were acquired at the baseline.

Table 1 summarizes the demographics of the subjects. This study was approved by the Institutional Review Board at the National Center of Neurology and Psychiatry, Tokyo, Japan.

**Table 1.** Characteristics of the study participants

|  | Normal data base |  | Evaluation data set |  |  |
| --- | --- | --- | --- | --- | --- |
| Source | NCNP | J-ADNI | J-ADNI |  |  |
| Filed strength | 3 T | 1.5T | 1.5 T |  |  |
| Category | NC | NC | sMCI | pMCI | AD |
| Number of participants | 1089 | 146 | 102 | 112 | 147 |
| Mean Age (*SD*) | 58.5 (14.3) | 67.6(5.66) | 72.8 (6.10) | 73.0 (5.54) | 73.5 (6.60) |
| Male (*Female*) | 375 (714) | 68(78) | 57(45) | 47(65) | 63(84) |

### Segment and calculate brain structure volume by multi-atlas fusion

We used the segmentation procedure incorporates a joint-label fusion method [6,10] and corrective learning [11], with the automated selection of thirty atlases. Each atlas included 131 manually traced ROIs and T1-weighted images. Briefly, the segmentation procedure involves the following algorithms: 1) The target T1-weighted brain image was separated to 30 large-regions by non-linear warping from the large-regions atlas in MNI space to the target T1-weighted image. The large-regions atlas created by grouping some ROIs to large region in the “neuromorphometrics atlas” attached CAT12 toolbox [1] based on the anatomical prior knowledge. 2) The 5 atlases were automatically selected from 30 manual traced atlases, according to the Pearson correlation between 30 atlases and the target image for each large-regions. 3) We create DARTEL template from the 5 selected atlases and the target image at each large-regions. Subsequently, 5 atlas were warped to target image via the large-region DARTEL templates. 4) Using the joint label fusion (JLF) method and SegAdapter, target image was segmented to 133 ROIs fused selected atlas at each large-regions; 5) The volumes of the ROIs are calculated after considering partial volume effects by the posterior probability maps based on the CAT12 toolbox. Since the segmentation procedure can perform parallel processing for each large-regions, so it led to a reduction in processing time.

The segmentation procedure was executed using a multi-core computer system (OS: CentOS 7.2, CPU: Intel Xeon E5-2600 24 cores, memory: 96GB), and the processing time was measured.

### Calculating Process for harmonized Z score

According to the report of Ma et al. [7], we can model and harmonize the undesired covariates, i.e. age, field strength, eTIV and sex, by general linear model. We defined the structure volume and all the other covariates as

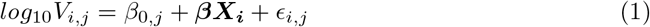

where *V_i,j_* is structural volume for i-th subject and j-th ROI and **X_i_** are design matrix of covariates, i.e. age, filed strength (0:3 T, 1:1.5 T), gender (0:female, 1:male) and *log*_10_(eTIV). After removing the effects of covariates, we can calculate the harmonized Z-scores at each ROIs as

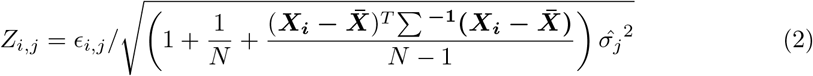

where 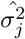 is unbiased distribution, *N* is number of participants and ∑ is covariance matrix of subjects.

We calculated the harmonized Z scores for each ROIs from 1089 NC subjects scanned by 3T MRI at NCNP and 146 NC subjects by 1.5 T provided from J-ADNI data set. Moreover, we evaluated and excluded the outliers of Z scores used Smirnov-Grubbs test with the 5% significance level, subsequently we recalculated z scores according to the same process on python 3.6.6.

In this article, we selected and presented some covariate combinations to illustrate scatter plots of age correlations : (a) no harmonization (raw variable); (b) field strength only; (c) the combination of field strength and eTIV;(d) the combination of field strength, eTIV and sex. Therefor, to calculate harmonized Z-score considering four situations: (a) age only; (b) the combination age and field strength; (c) the combination of age field strength and eTIV; (d) the combination of age, field strength, eTIV and sex.

### Evaluation of harmonization of different filed strength and sex

We used the Kolmogorov-Smirnov test to measure the separation of the z-scores distributions and quantitatively compare the separation of the sample distributions compare the Z-scores between NC from 1.5T and 3 T scanner. If harmonization well works, the distance of Z-score distribution in subgroups with different covariates (i.e. filed strangles and sex) in NC subjects well be smaller.

### Evaluation of harmonized z-score for AD detection and classification sMCI/pMCI

Finally, we evaluated the detectability of the harmonized Z-score in the ROIs between AD and NC group, and sMCI and pMCI group. The harmonized Z-score was analyzed by receiver operating characteristic (ROC) curves and the area under the curve (AUC). The ROC curves of each ROI were drawn based on the trade-off between sensitivity and specificity for discriminating diagnostic group, and higher AUC indicates better detectability by the harmonized Z-score in the ROIs.

## Results

We perform the segmentation procedure all T1 brain MRI, most of the processing was completed within 30 minutes.

### Evaluation of harmonization of different filed strength and sex

Fig. 1 shows the correlation between raw and harmonized volumes of a set of 8 ROIs associate with cognitive function for the NC group. The male/female and 1.5 T/3 T data are shown overlapped. The blue line represents the regression line for age, and the red and green lines are 95% confidence intervals. An overall negative correlation between ROI volumes within gray matter and age is found. The coefficients for harmonized Z-score calculation in all ROIs are listed in the Appendix data.

**Figure 1.**
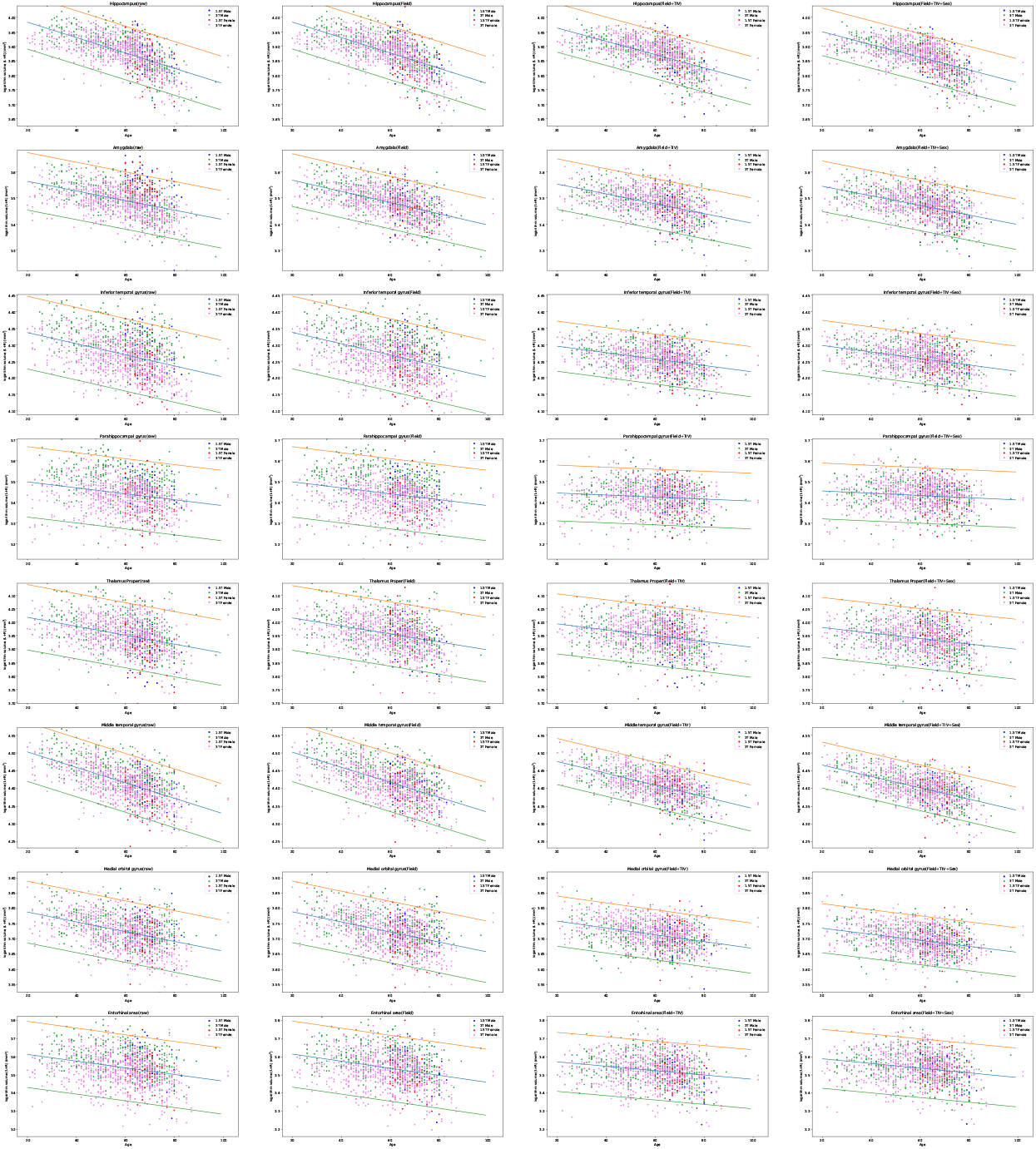
Correlation between the age (x axis) and some selected structure logarithm volumes (y axis) for the NC group for 1.5 T males (blue), 3 T males (green), 1.5 T females (violet) and 3 T females (red).

The 95% confidence intervals became narrow after each harmonization, and were most improved when the magnetic field strength and eTIV or all of covariate were corrected. To quantitatively assess the shift of kernel density estimate function of Z-score before and after harmonization accounting for each covariate, we performed the K-S test between 1.5T NC and 3 T NC group. Before harmonization, in 72/133 ROIs the null hypothesis that the two sample structural volume distributions scanned by 1.5 T MRI and 3 T MRI come from the same population were rejected, however not rejected after harmonization.

Fig. 2 shows the kernel density estimation function (KDEF) of the harmonized Z-score. The KDEFs of Z-score showed the brain atrophy of AD on these structures compared to the NC group. When the Z score shows negative value, particularly less than −2, it indicates that the ROI of subjects is significantly atrophied compared to the NC. After harmonization, the distribution of Z-scores in AD and pMCI group were shifted to leftward at hippocampus, amygdara, infeiror temporal gyrus, para hippocammpal gyrus middle-temporal gyrus, this things shows that the detection power of AD by Z-score is improved, after harmonization including all covariates, filed strength, eTIV and sex.

**Figure 2.**
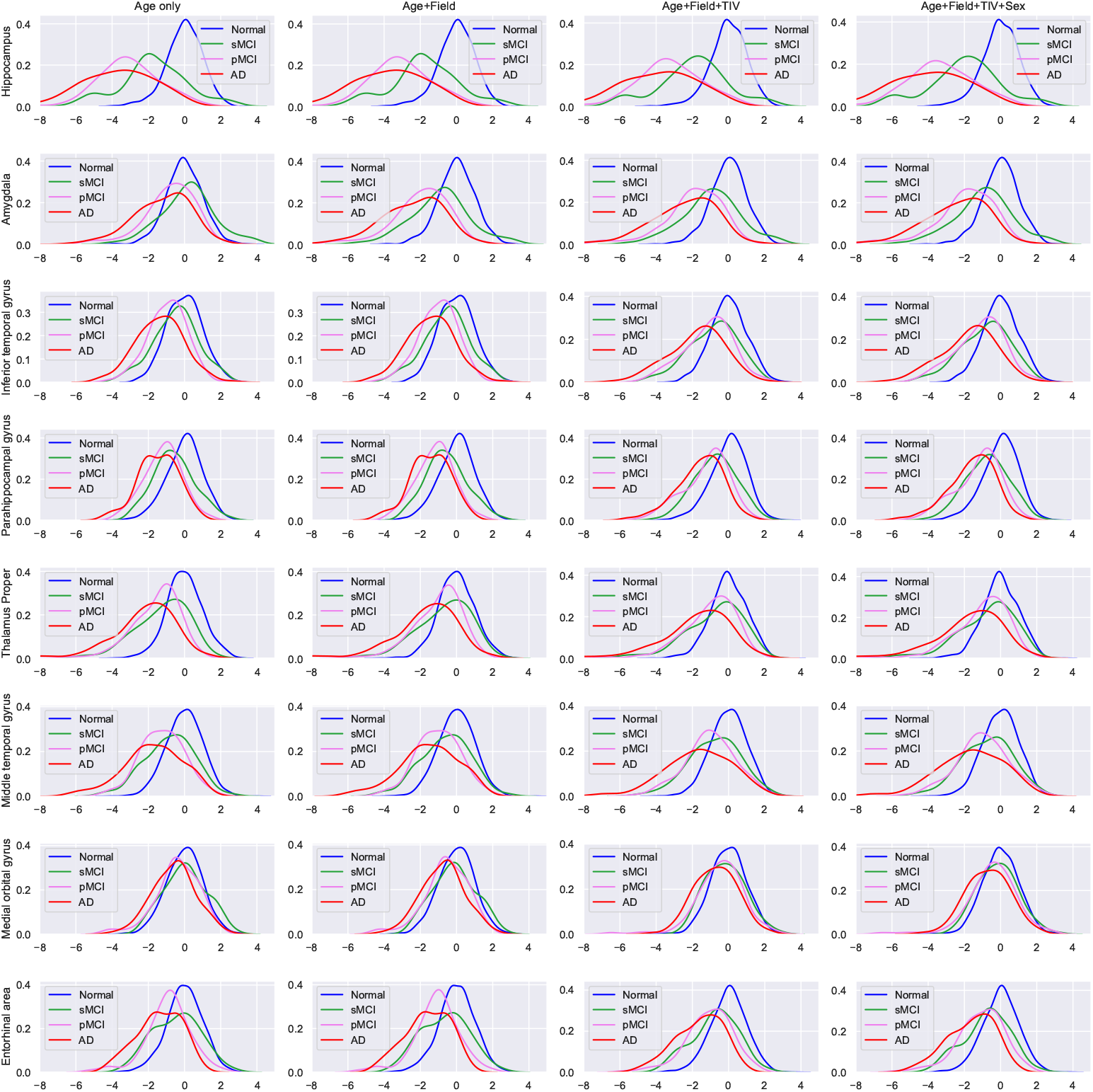
The kernel density estimation function of the harmonized Z-score taken from a select few gray matter structures. Blue line=NC, green line=sMCI, violet line=pMCI, red line=AD

### Discriminative power between NC vs. AD and sMCI vs. pMCI

The AUC values compared with AD vs. NC and pMCI vs sMCI are shown in Table 2. The results showed that the Z-score of the lateral hippocampus temporal showed high AUC values, right hippocampus is 0.96 and left is 0.95.

**Table 2.** The area showed the AUC value of 80 or more for NC vs. AD, and of 60 or more for sMCI vs. pMCI.

| AD vs NC |  | pMCI vs sMCI |  |
| --- | --- | --- | --- |
| Region | AUC | Region | AUC |
| Right Hippocampus | 0.96 | Right Hippocampus | 0.70 |
| Left Hippocampus | 0.95 | Left Hippocampus | 0.68 |
| Right Amygdala | 0.88 | Left Amygdala | 0.66 |
| Left Amygdala | 0.86 | Right Amygdala | 0.64 |
| Right ITG | 0.82 | Right MOrG | 0.60 |
| Left Thalamus Proper | 0.80 | Right Entorhinal area | 0.60 |
| Left PHG | 0.80 | Right Amygdala | 0.60 |
| Right PHG | 0.80 |  |  |
| Right MTG | 0.80 |  |  |

PHG, Para Hippocampal Gyrus.
ITG, Inferior Temporal Gyrus.
MTG, Middle Temporal Gyrus.
MOrG, Medial Orbital Gyrus.

## Discussion

We analysed 131 brain structural volume of 1235 (3T=1089, 1.5T=146) cognitive normal subjects and it allowed the calculation of harmonized Z-score considering age, filed strength, eTIV and sex. The harmonization of theses covariate improved the reliability of normal database because confidence interval became narrow after harmonization. Quantitative evaluation based on Kolmogorov-Smirnov test showed that the null hypothesis of the two sample structural volume distributions scanned by 1.5 T MRI and 3 T MRI come from the same population was not rejected. This results showed that the harmonization worked well.

Out method achieved the high AUC values (0.95) for discrimination NC and AD, the moderate AUC values (0.70) for classification of pMCI and sMCI. Elahifasaee et al archived the classification accuracy of 65.94% can be achieved for classification of pMCI and sMCI based on feature decomposition and kernel discriminant analysis [3]. Several studies have shown that the discrimination between sMCI and pMCI is difficult task. Our results can not possible direct comparison Farzaneh’s result because the metrics are different, but considering that our results based on only univariate Z-score, it can be said that our results showed good accuracy to classify sMCI and pMCI.

From these results, our method can be effective not only for AD detection with high accuracy but also for sMCI/pMCI discrimination without affect of the different field strength, in other words high generalization capability, because most of our database consists of 3T MRI subjects and the evaluation data set derived from 1.5T MRI subjects. Furthermore, our method can be expected to improve the accuracy by the multivariate Z-score approaches, such as machine learning and other multivariate analyses. We believe that multivariate Z-scores derived from whole brain ROIs can be applied to the type-classification of the dementia, and the biomaker of another neurodegenerative disorders.

## Acknowledgments

We would like to offer my special thanks to Nippon Tect Systems Co. Ltd. for providing free version of the segmentation procedure.

